# DAtest: a framework for choosing differential abundance or expression method

**DOI:** 10.1101/241802

**Authors:** Jakob Russel, Jonathan Thorsen, Asker D. Brejnrod, Hans Bisgaard, Søren J. Sørensen, Mette Burmølle

## Abstract

DAtest is an R package for directly comparing different statistical methods for differential abundance and expression analysis on a dataset of interest; be it data from RNA-seq, proteomics, metabolomics or a microbial marker-gene survey. A myriad of statistical methods exists for conducting these analyses, and with this tool we give the analyst an empirical foundation for choosing a method suitable for a specific dataset. The package supports categorical and quantitative variables, paired/block experimental designs, and the inclusion of covariates. It is freely available at GitHub: https://github.com/Russel88/DAtest along with detailed instructions.

## Introduction

Identifying dataset features that are associated with variables of interest, such as design groups, technical or environmental covariates, is a common procedure in the analysis of microbial marker-gene (e.g. 16S rRNA amplicons), RNA-seq, metabolomics and proteomics data. There are no gold standards for these types of analyses, but the choice of method can change results considerably, concerning the rate of false positives and the power to detect a signal^1–3^.

Here we introduce DAtest, an R package for comparing differential abundance and expression methods on a specific dataset. We embrace that no methods work well on all datasets, and the aim is therefore to aid the analysts in choosing an appropriate method for their dataset, such that the choice of statistical method is well-grounded. The package provides a simple and reproducible framework for estimating false positive rates (FPR) and Area Under the Receiver Operating Characteristic (ROC) Curves (AUC) of 25 different methods (38 including variations of similar methods), and it supports categorical and quantitative variables, paired/block experimental designs, and the inclusion of covariates. It runs in parallel for fast performance, it is platform independent, and the software, including tutorial, is freely available at GitHub (GPL 3.0): https://github.com/Russel88/DAtest.

## Methods

Common to all ‘omics datasets is that they usually have many more features than samples, the large *p*, small *n* problem, and the main difficulty in analyzing these datasets is controlling the false positive rate while maintaining statistical power.

The main function, testDA, estimates the FPR and AUC for each method applied to a given dataset. The procedure goes as follows:

1. Shuffle predictor variable (E.g. case vs. control)
2. Spike in data for some randomly chosen features such that they are associated with the shuffled predictor
3. Apply all relevant methods
4. Construct confusion matrix and estimate FPR for each method
5. Calculate AUC for each method
6. Repeat (1) to (5) several times for reliable estimations

Below is a detailed description of each step in this procedure:

First, the predictor variable is shuffled. If the experimental design contains blocking, samples are paired, or there are repeated measures, the predictor is shuffled within the blocks or subjects. Covariates are left untouched.

Second, randomly chosen features are spiked with a set effect size. If the predictor is categorical, one class, from the shuffled predictor, is spiked by multiplying the original values with the effect size (script adapted from ^1^). If the predictor is quantitative the original values are spiked as follows:

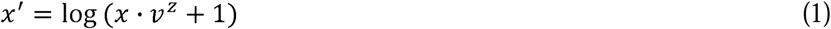

Where *x* are the original values, *x’* the spiked values, *v* the effect size (from zero to infinity, 1 is no effect), and *z* the z-scaled shuffled quantitative predictor variable. *x’* is then rescaled such that the range of values is similar to *x*. If the dataset is compositional, e.g. for RNA-seq or OTU data, the dataset is renormalized such that the library sizes are unchanged after spike-in.

Third, all relevant methods are applied. Included are classical tests such as t-test, Wilcoxon rank-sum test, (generalized) linear regressions (including zero-inflated versions); tests originally developed for microarray data but adapted to sequence data, such as limma^4^ and SAMseq^5^; widely used methods for RNA-seq data, such as edgeR^6^ and DESeq2^7^; and methods dedicated for microbiome data such as metagenomeSeq^8^ and ANCOM^9^. A permutation test for paired data was developed for this package: It is similar to the one from Thorsen et al. (2017)^1^, except that the predictor is shuffled within each pair, and the fold change is calculated for each pair before averaging. It is also possible to add multiple user-defined methods. See Supplemental Table for details on implemented methods.

Fourth, as we know which features are spiked, a confusion matrix can be constructed and the FPR estimated. If a large number of features are compared to a randomly generated variable, we should expect 5% of the nominal p-values to be lower than or equal to 0.05 due to the uniform distribution of p-values under the null hypothesis. Therefore, we expect a 5% FPR from shuffling the predictor variable.

Fifth, for estimating the statistical power of the methods we use the area under an ROC curve constructed from the nominal p-values. For a perfect test, ranking the p-values should provide complete separation of the spiked and non-spiked features and AUC = 1. In practice, it will be anywhere between 0.5 and 1, where 0.5 corresponds to no separation. For ANCOM and SAMseq, that do not output p-values, the ROC curve is constructed from the test statistics as these provide complete separation of the detected and non-detected features. Finally, methods can be compared with the generic R functions plot and summary. For the final analysis with the original data, all methods are easily run from the same standardized input used for the testDA function. Alternatively, many methods can be run simultaneously and compared e.g. with Venn diagrams.

## Results

We tested our package on data from the Human Microbiome Project^10^ (https://www.hmpdacc.org/hmp/HMQCP/). To illustrate the usefulness of the package we demonstrate how including information on pairing of the samples influence the performance of the different methods. Two subsets are used: 20 pairs of samples from “Subgingival plaque” and “Supragingival plaque”, and 50 pairs of samples from “Throat” and “Tongue dorsum”. All samples are from first visit, and only from patients that have both samples. OTUs that are present in less than 4 samples and/or have less than 100 total reads are grouped in a single feature named “Others” (See Supplemental Script for details). In general, the methods perform better with paired tests, with higher AUC and lower FPR (Fig. 1). Importantly, edgeR (erq and erq2) and DESeq2 (ds2x), which both have high AUCs, can only adequately control the FPR with pairing. Furthermore, Wilcoxon Signed-Rank test (wil, paired) has considerably higher sensitivity than Wilcoxon Rank-Sum test (wil, unpaired).

**Fig. 1.**
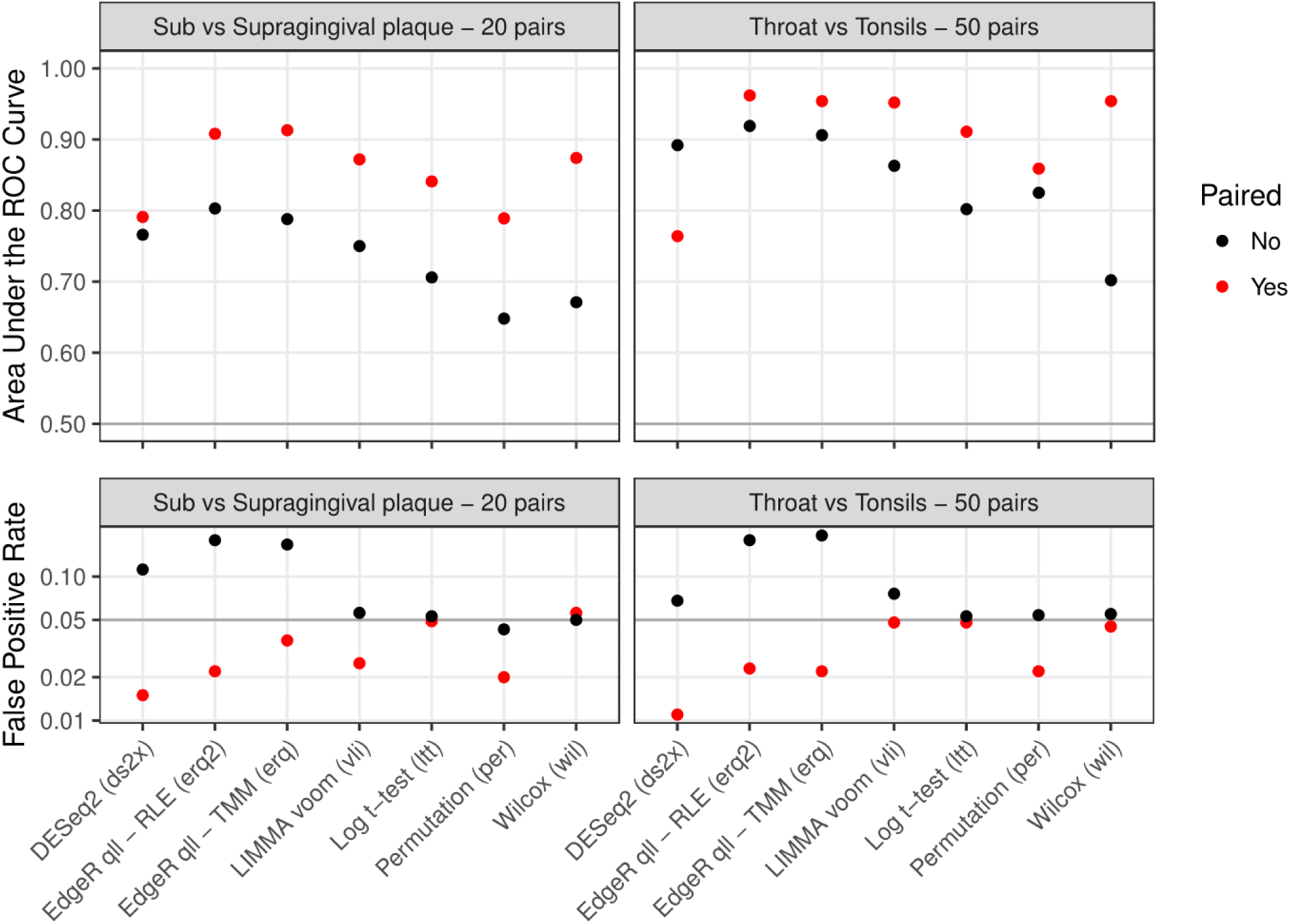
AUC and FPR can change dramatically depending on whether the statistical model uses information on the pairing of the samples. Points are medians from 10 runs with the testDA function. The same OTUs were spiked for the paired and unpaired tests. RLE: Relative Log Expression normalization, TMM: Trimmed Mean by M-value normalization.

## Discussion

Benchmarks has shown how sparsity and sample size considerably change the performance of differential abundance methods (see e.g. ^1^). Here we show that including information on pairing of the samples also change performance of the methods, mostly for the better. Choosing the correct differential abundance or expression method is difficult, since many factors influence the performance of the methods. Here we present a tool that aids the analyst in choosing a method, based on a simple, efficient, reproducible and statistical sound methodology that is based on the unique characteristics of his/her own dataset. As the DAtest R package is open source and easily extendable, it provides a framework for further methodological development.

## Author Contributions

J.R. developed and maintains the software. J.T., and A.D.B. contributed to the development. H.B., S.J.S., and M.B. supervised the development. J.R. wrote the manuscript. All authors commented on the manuscript.

